# Tuning methylation-dependent silencing dynamics by synthetic modulation of CpG density

**DOI:** 10.1101/2023.05.30.542205

**Authors:** Yitong Ma, Mark W. Budde, Junqin Zhu, Michael B. Elowitz

**Affiliations:** Division of Biology and Biological Engineering, California Institute of Technology, Pasadena, CA 91125, USA; Primordium Labs, Arcadia, CA 91006, USA; Department of Biology, Stanford University, Stanford, CA 94305, USA; Howard Hughes Medical Institute, California Institute of Technology, Pasadena, CA 91125, USA

**Keywords:** KEYWORD: Epigenetics, DNA methylation, DNMT3b, Synthetic biology

## Abstract

Methylation of cytosines in CG dinucleotides (CpGs) within promoters has been shown to lead to gene silencing in mammals in natural contexts. Recently, engineered recruitment of methyltransferases (DNMTs) at specific loci was shown to be sufficient to silence synthetic and endogenous gene expression through this mechanism. A critical parameter for DNA methylation-based silencing is the distribution of CpGs within the target promoter. However, how the number or density of CpGs in the target promoter affects the dynamics of silencing by DNMT recruitment has remained unclear. Here we constructed a library of promoters with systematically varying CpG content, and analyzed the rate of silencing in response to recruitment of DNMT. We observed a tight correlation between silencing rate and CpG content. Further, methylation-specific analysis revealed a constant accumulation rate of methylation at the promoter after DNMT recruitment. We identified a single CpG site between TATA box and transcription start site (TSS) that accounted for a substantial part of the difference in silencing rates between promoters with differing CpG content, indicating that certain residues play disproportionate roles in controlling silencing. Together, these results provide a library of promoters for synthetic epigenetic and gene regulation applications, as well as insights into the regulatory link between CpG content and silencing rate.

## INTRODUCTION

Methylation of CG dinucleotides (CpGs) plays critical roles in mammalian development, tumor progression, and aging.^1–4^ These functions result mainly from the ability of CpG methylation to induce and stabilize gene silencing in mammals through multiple mechanisms.^5, 6^ Control of DNA methylation and further gene silencing depends on both trans-acting factors and the DNA sequence itself. Trans-factors include methylation “writers” such as DNA methyl-transferases (DNMTs)^7^, and “erasers” such as TET1^8^ that establish and alter methylation marks, as well as “readers”, such as MeCP2 and histone deacetylases^9^ that link methylation to regulation of gene transcription.^6, 10^ In mammalian cells, DNA methylation occurs mainly at CpGs. As a result, the distribution of CpG dinucleotides within a given sequence can play a pivotal role in methylation-based gene regulation.

At the genome level, regions with different CpG content exhibit distinct methylation patterns,^11, 12^ potentially due to cooperativity between nearby CpGs and other suppressive epigenetic marks, which can generate positive feedback.^13, 14^ Relatively high CpG-density regions (CpG islands) from a human chromosome largely maintain their methylation state when hosted in a transchromosomic mouse model,^15^ suggesting that DNA sequence composition plays a strong role in establishing stable methylation states. Conversely, insertion of several hundred base pairs of CpG-free DNA can disrupt these patterns, permitting de novo methylation of the surrounding CpG island.^16^ However, the precise role of CpG sequence context can be difficult to discern at natural loci, where regulation is also affected by many other cis- and trans-acting factors, including cell-type specific methylation writer and reader profiles, neighboring (non-CpG) motifs that recruit epigenetic modifiers, pre-existing chromatin states, etc. To clearly delineate the role of DNA sequence in methylation and silencing, one would ideally want to directly compare the methylation and silencing of similar promoters with different CpG distributions, in the same genomic context. Further, because methylation and its effects on gene regulation can both be dynamic ^17–19^, the ability to control the timing of methyltransferases (DNMTs) recruitment to a locus and follow the resulting changes in gene expression is also desirable.

The field of synthetic epigenetics seeks to harness epigenetic regulatory mechanisms to control gene expression on different timescales.^20, 21^ Recent work demonstrated the ability to regulate synthetic or endogenous gene expression by recruiting DNMTs to specific target genes,^18, 22–24^ and even create fully synthetic DNA methylation-based systems for synthetic epigenetic memory.^25^ In CHO-K1 cells, transient DNMT recruitment to a locus drives stochastic, irreversible, all-or-none silencing over timescales of about 5 days.^18^ However, it is unclear how the dynamics of gene silencing depends on the DNA sequence of the regulated target gene. Understanding the effects of sequence composition on silencing rates could provide insight into gene regulation by DNA methylation and also expand the synthetic epigenetic toolbox for fine tuning circuits.

Here, we adapted a previously established DNMT recruitment system to analyze the effects of DNA sequence on methylation-dependent silencing.^18^ We derived a library of promoters with different CpG densities from a synthetic promoter and observed the relationship between CpG density and the silencing dynamics occurring after DNMT recruitment. Using a mathematical model of methylation, together with sequencing identifying methylation marks, we showed that the observed gene expression dynamics could be explained by methylation accumulation on the DNA. We also identified several specific CpG elements that appear to play disproportionate roles in silencing dynamics and confirmed that one of them (near the TATA box and transcription start site), causes significant changes in silencing dynamics. Our results reveal how CpG density influences silencing dynamics, and provide a library of promoters with different silencing rates for synthetic applications.

## RESULTS AND DISCUSSION

### Construction of a promoter library with varying CpG content

To investigate the relation between promoter CpG content, and its DNMT-dependent silencing rate, we adopted a previously described synthetic methylation-silencing system.^18^ In this system, the catalytic domain of DNMT3b (DNMT3bCD) is fused with a reversed tetracycline repressor (rTetR), allowing precise temporal control of the recruitment to a target gene by adding doxycycline (dox) to the culture media.^26^ Here, to focus on the role of DNA methylation in silencing, we specifically used the mouse gene Dnmt3b’s catalytic domain, spanning amino acids 402-872. This region omits the major heterochromatin-interacting PWWP domain.^27^ This construct also incorporates a co-expressed H2B-mCherry fluorescent protein fusion. We stably integrated this construct using the piggyBac transposon system, and sorted cell populations with similar mCherry fluorescence level (Figure S1A). This enabled direct comparison of different promoters (see below) with the same DNMT3bCD expression context.

As the target gene, we used an H2B-mCitrine^28^ fluorescent fusion protein. This target was driven by one of a set of promoters containing varying densities of CpG (see below). In each promoter, an array of 5 rTetR binding sites (TetO) was fused upstream of the promoter, allowing recruitment of rTetR-DNMT3bCD (Figure 1A). To enable direct comparison between target promoters at the same genomic context, all reporter cassettes were site-specifically integrated as a single copy into the epigenetically active φC31attP/attB landing site within an artificial chromosome that was previously engineered into CHO cells (MI-HAC CHO cells, also see MATERIALS AND METHODS).^29^

**Figure 1.**
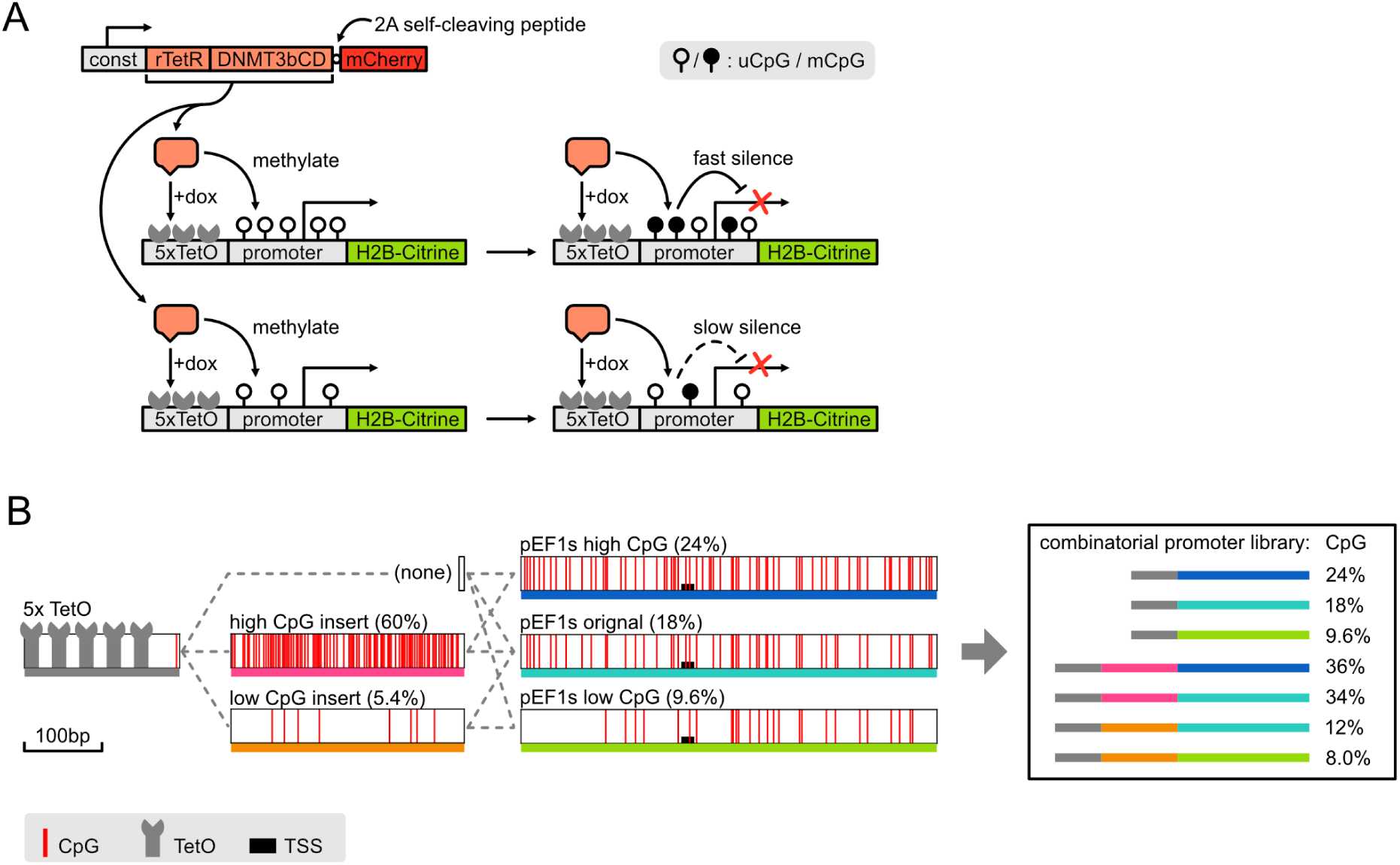
rTetR fused DNMT3b catalytic domain methylates and silences a reporter library of different CpG content upon recruitment: A. Schematic of the synthetic methylation-silencing system. rTetR fused DNMT3bCD is expressed constitutively, and upon induction of dox, recruited to the promoter region of a site-specifically integrated Citrine reporter. The recruitment methylates the promoter and further silences the gene expression, with dynamics depending on the promoter’s CpG content. B. Design of the library of promoters with varying CpG content. 5x tandem TetO binding sites were fused with an insert (or no insert) and a pEF1s synthetic promoter to make the library of promoters. Red lines represent CpG dinucleotides, and the pEF1s promoters are vertically aligned to show sequence homology.

We constructed a library of synthetic promoters that differed in their CpG densities. We started with a synthetic version of the human elongation factor 1α promoter (pEF1s(orig), with 18% CpG density), a 544bp fusion of promoter fragments from the human EF1α promoter and human T-cell virus (HTLV),^30^ that has been commercially available (InvivoGen) and has been used for antibody expression and gene therapies.^31, 32^ To identify conserved CpG elements, we compared both the EF1α fragment and HTLV fragments of this promoter to their natural orthologs respectively.^33^ We then removed or added CG pairs into the promoter at non-conserved sites. With this procedure, we generated promoters with varying CpG densities at 9.6% and 24% (pEF1s(low) and pEF1s(high) respectively, Figure 1B, middle).

Next, to broaden the range of CpG densities, we designed an additional DNA segment, inserted upstream of the promoter, containing high (60%) CpG density (Figure 1B, high CpG insert). We altered this CpG insert by swapping out CG with GC dinucleotides, or by replacing C with T, to create a lower CpG density (5.4%) insert, while otherwise preserving its sequence similarity with the high CpG insert (Figure 1B, low CpG insert). Altogether, we combined the three pEF1s promoters with the two inserts, or with no insert, to produce a library of 7 sequences whose overall CpG density ranged from 8.0% to 36% (Figure 1B right). Despite their variation in CpG density, all 7 promoters drove strong expression of the fluorescent protein reporter, producing ∼200-fold greater signal compared to autofluorescence in non-transfected cells, with a 2.8 fold variation of expression level (Supplementary Figure S1B and S1C). This difference could be either due to the difference in the promoter sequences, or due to altered mRNA secondary structure of 5’UTR (part of the altered promoter sequence), resulting in changes in RNA half-life^34^. However, this expression change did not impact quantification of the silenced fraction, as silencing thresholds were set independently for each promoter (see below). As expected, the original pEF1s promoter shows the highest activity, while our alterations to the sequence (both addition or reduction of CpGs) slightly lowered its activity. Therefore, there was no correlation between expression level and CpG content.

### DNMT-dependent silencing rate correlates with promoter CpG density

Previous analysis of silencing dynamics by DNA methylation in a similar system revealed that transcriptional silencing occurs through stochastic, all-or-none, irreversible events in individual cells.^18^ To confirm that our system has similar kinetics, we induced DNMT3bCD recruitment to promoters for 4 days and then released the recruitment for 2, 6 and 10 days, and measured the Citrine fluorescence via flow cytometry for all three promoter variants (Figure 2A, Supplementary Figure S2A and S2B for pEF1s(high), pEF1s(orig), and pEF1s(low), respectively). As expected, a fraction of the cells were transcriptionally silenced after the induction, and gradually diluted out stable H2B-Citrine fluorescent protein during the “release” phase due to cell divisions (approximately 22 hours per division, observed by the shifting position of silenced population peaks). Meanwhile, the active populations (peaks on the right) remained stable in terms of both fluorescence level and cell population fraction, indicating an all-or-none, irreversible kinetics as reported before.

**Figure 2.**
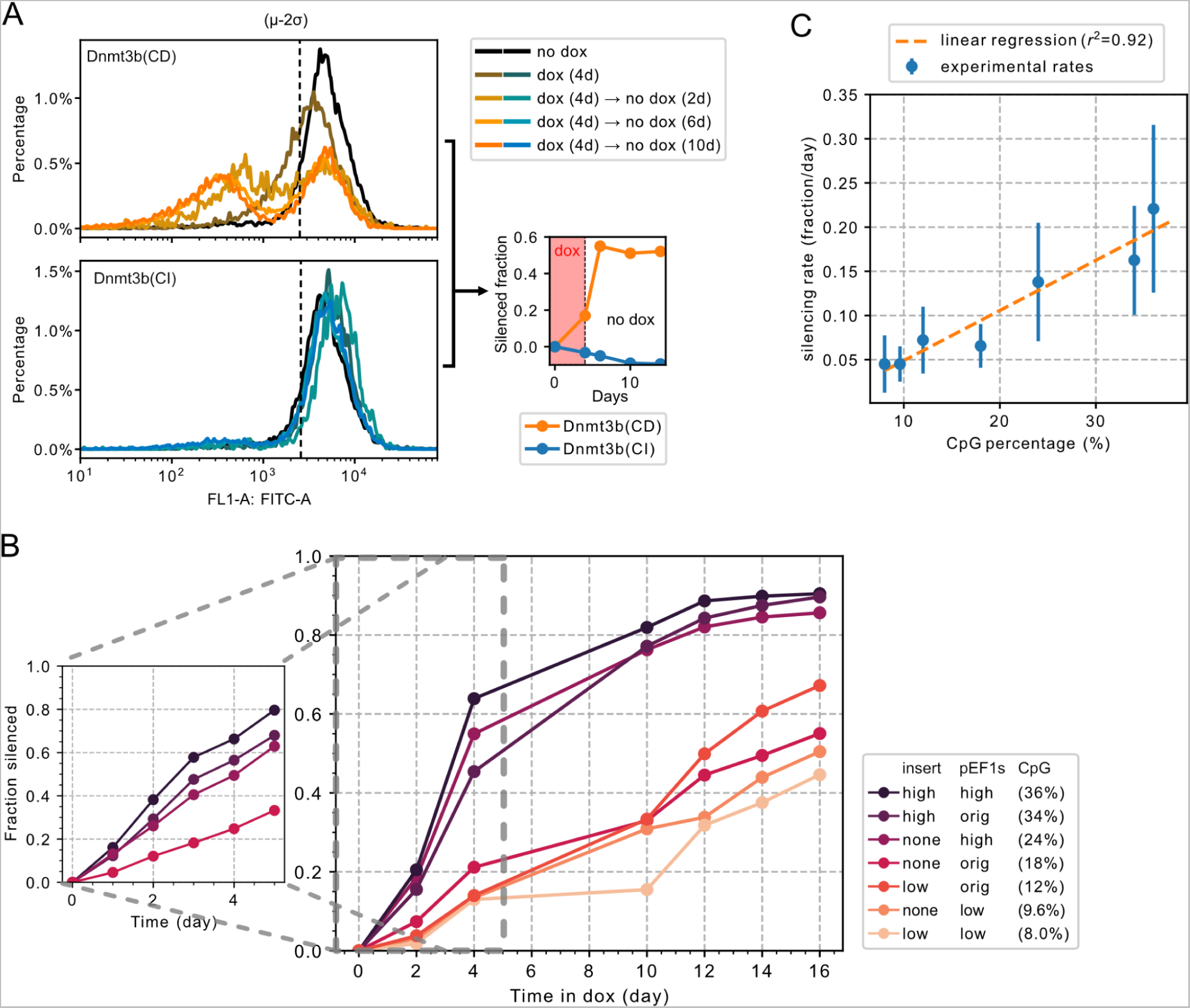
Promoters’ silencing rate correlates with their CpG content. A. Promoter silences with all-or-none kinetics when DNMT3bCD is recruited to the locus, and this silencing is dependent on DNMT3b’s catalytic activity. Cells with the pEF1s(high) promoter were treated with dox for 4 days, and then no dox (release) for 2, 6, and 10 days. Cells are analyzed by flow cytometry at various time points (left), and quantified by comparing to the no dox control (black lines in the left). A log-normal distribution is first fitted onto the no dox control’s positive population, and μ-2σ are used for quantification of silenced fractions. These fractions are then normalized to no dox controls’ silenced fraction (see MATERIAL AND METHODS, right). The silenced fractions are stable after the release of dox for 2 days, and no silencing is observed in the DNMT3bCI controls. B. Time course of the silenced fraction of different promoters. Cells were treated with or without dox, and then with 2 days of no dox (release). The fraction of silencing is determined as described in A: cells with lower fluorescence than μ-2σ of the no dox control group were determined as silenced. The silencing rate is further normalized to the no dox group (see MATERIAL AND METHODS). For the shorter time scale (left), the same method is used except with a higher time resolution. C. Summary of the silencing rates in B. The silencing rate is calculated by subtraction between each pair of neighboring dots, and then normalized by time intervals in between, as well as the remaining fraction size. We omitted dot pairs over 80% fraction as the normalization fraction is too small.

Setting a silencing threshold 2 standard deviations (2σ) below the mean fluorescence levels of actively expressing (“no dox”) cells yielded a stable value after 2 days of release (see quantification on the right in Figure 2A, Supplementary Figure S2A and S2B), allowing consistent quantitation of all-or-none silencing. Because the control groups were single peaked and exhibited consistent variation, this cutoff occurred at values ranging from 54% to 71% of the mean, depending on cell line (Figure S2C).

We also used a catalytically inactive version of the DNMT3bCD protein (P656V and C657D double point mutations,^35^ noted as DNMT3bCI) as a negative control. Even though the CI version were expressed at a similar level of the CD version (Figure S1A lower half), no silencing effect was observed when they were recruited to the promoters (Figure 2A, Supplementary Figure S2A and S2B, lower half as well as blue lines in the quantifications). This confirmed that the observed silencing effects resulted specifically from DNA methylation activity, and not other protein interactions or interference with transcription machinery.

These results established that the promoters silence in a methylation-dependent, all-or-none and irreversible manner, and indicate that silencing kinetics can be captured by the dox induce-and-release protocol.

To quantify silencing dynamics across the library, we analyzed the dynamics of silent cell accumulation over a time course (up to 19 days) of dox induction with 2 day subsequent release of the recruitment at various time points (Figure 2B, right). As some of the promoters, notably those with insert(high) or pEF1s(high), reached over 50% of silencing only after four days (about two time points in our setup), we added a biological replicate for each of the four fastest promoters with a separate time course with finer time resolution (Figure 2B, left). From the time-course, we can conclude that higher overall CpG density on the promoter results in faster silencing dynamics. This result is robust to analysis with a stricter cutoff of 90% expression reduction (Figure S2B). Even with the less sensitive threshold, we observed a similar trend of increased CpG density correlating with faster silencing.

The time-course dynamics for each promoter could be summarized by an empirical silencing rate as the silenced fraction per unit time (day), normalized by the remaining active population. Silencing rates varied over nearly an order of magnitude across promoters with various CpG densities. Further, the silencing rate correlated linearly with CpG density over this range (Figure 2C). These results show that across varying CpG densities, DNMT-dependent silencing dynamics are broadly consistent with a stochastic, all-or-none silencing process, occurring at a rate that depends on CpG density.

### Silencing kinetics follow a stochastic switching model

Previous studies using a similar system with a different version of the EF1α promoter showed that silencing kinetics could be described as a single-step stochastic switching event from the active to the silent state^18^ (Figure S3A). In the model, each promoter silences stochastically at a time-invariant rate *β*. Here we tested whether a similar model could fit our data, if we allowed *β*(*c*) to depend on CpG density, . With this assumption, the size of the active population fraction, *c*, can be described by a simple differential equation:

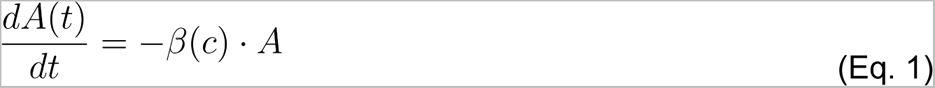

Note that cell proliferation does not need to be explicitly incorporated due to the heritability of the expression state. In this model, the active population, *A*, decays exponentially over time *t*, . To test this model, we plotted the time course data (Figure 2B) in terms of the remaining active population, *A*(*t*) (Figure S3B), and observed a linear-log relationship, consistent with exponential decay, for every promoter variant. In these plots, the switching rate *β*(*c*) ranged from 0.032 d^-1^ to 0.274 d^-1^, depending on CpG density. These results are consistent with a simple stochastic switching model in which silencing rate is tuned by CpG density.

### Methylation accumulates after DNMT recruitment

Given that promoter silencing depends on the methylation activity of the recruited DNMT3bCD, we next asked whether methylation accumulates at similar or different rates for different promoters. We used FACS to isolate the transcriptionally active cell fraction (*A* in the model), at different times after dox addition (Figure 3A). We then measured promoter CpG methylation profiles using methylation-specific sequencing (EM-seq^36^) (Methods, Figure 3A).

**Figure 3.**
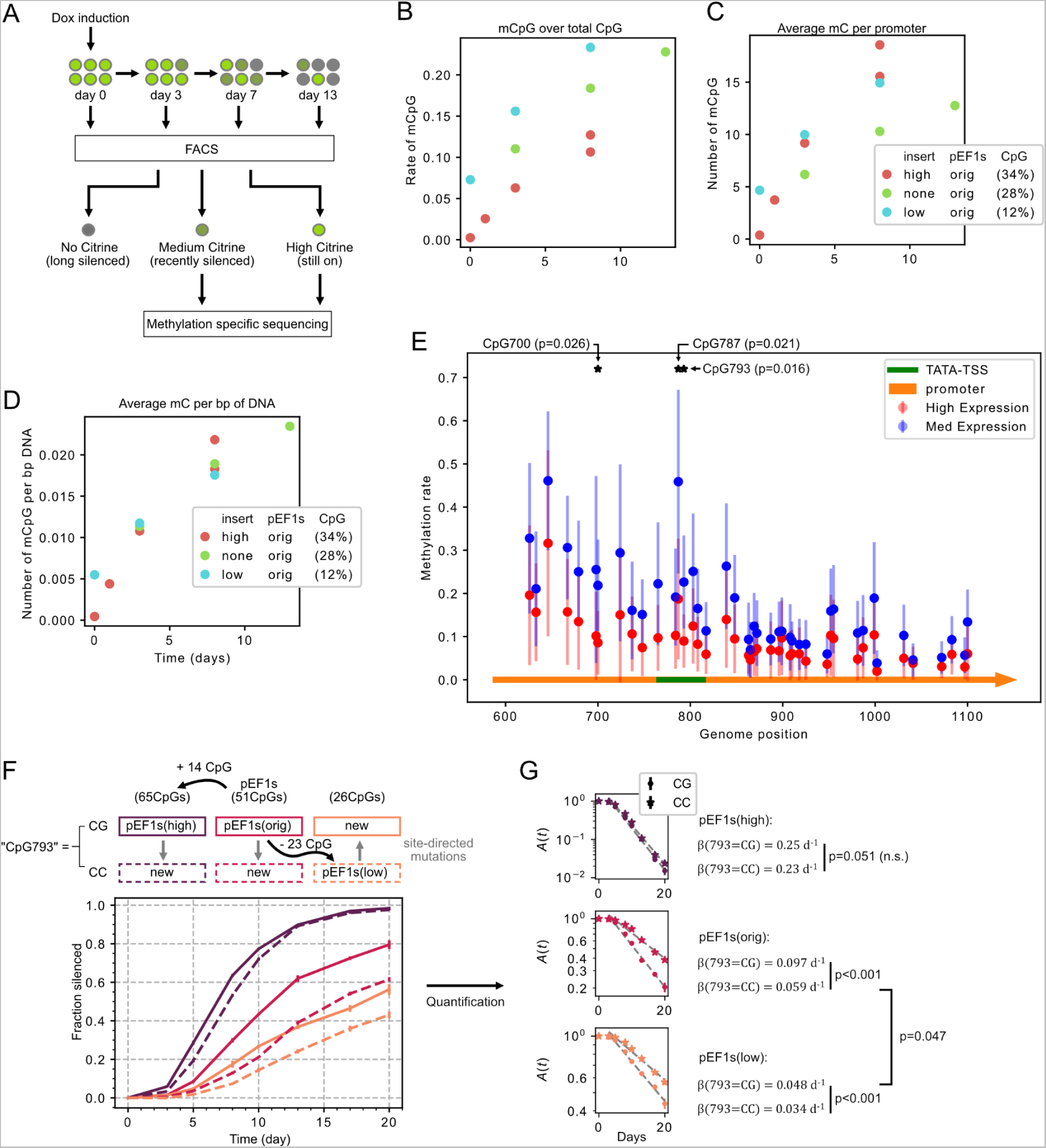
Sequencing reveals the constant accumulation of methylation, and potentially master CpGs. A. Schematics of the FACS-Sequencing experiment: Cells with different promoters are treated with dox, and then FACS-sorted to three bins based on the Citrine brightness (high, med and low), consisting cells that are “still ON”, “recently silenced” and “long silenced” respectively. The first two groups proceed to downstream methylation-specific sequencing (see MATERIALS AND METHODS). B-D. Total CpG methylation (B), methylation frequency (C) and total methylation normalized by promoter length (D) accumulates in the promoter with time in the “still ON” cell population. “Still ON” populations are sorted out as indicated in A at intended dates, and subsequently analyzed by methylation-specific sequencing (EM-seq), targeting the integrated gene promoter. E. CpGs around TATA-box and TSS (highlighted in green) show significant difference in methylation between the “still ON” and “recently silenced” group. Methylation percentages of different samples at different days were pooled together for comparison (a total of 10 from “still ON” group compared to 6 from “recently silenced” group). P-values are from student t-test. F. Mutation at CpG793 changes promoters’ silencing rate significantly. New cell lines are constructed by introducing point mutations (CG to CC or the inverse) at CpG793 in pEF1s(high), pEF1s(orig) and pEF1s(low) promoters (top). Cell lines are then constructed as described previously in this paper. DNMT3bCD recruitments are induced by dox at day zero and cells were analyzed by flow cytometry at each time point after dox induction. G. Quantification of the silencing rate (similar method as Supplementary Figure S3A and Eq. 1) of time course in F. We excluded the time point at 0 from the fitting as dox release was not included in this experiment. P-values are calculated based on the estimation and standard error or *β* from linear regression.

As expected, methylation accumulated in the transcriptionally active populations, as measured by methylation rate (methyl-CpG over total CpG), total methylation per promoter, as well as total methylation per promoter per bp of DNA (Figure 3B-3D respectively). Unexpectedly, however, the rate of methylation accumulation was independent of CpG density, measured as total methylation per bp of DNA (Figure 3D). In fact, the rate of methylation per CpG was greater at promoters with lower CpG densities (Figure 3B), while the total number of methylated CpG in the promoter region was similar across different promoters. This behavior is compatible with saturation of methylation capacity of the locally recruited DNMT3bCD. Alternatively, it could also reflect an effective interaction, in which unmethylated CpGs inhibit methylation at nearby CpG sites.^13^

### A single CpG has a disproportionate impact on silencing rate

The apparent discrepancy between the CpG density-dependent silencing rate and the density-independent methylation rate provoked the question of whether certain individual CpGs might play disproportionate roles in controlling silencing. Such CpGs would be expected to exhibit significant differences in methylation between cell populations containing active versus recently silenced promoters.

To discover such CpGs, we pooled all available sequencing results from different days. Within the pEF1s region, where all three promoters overlap (∼90% of the pEF1s region) (Figure 3E), we identified three CpGs with significantly different methylation levels between the two expression groups (p<0.05). Interestingly, two of the most significant CpGs, including the top ranked one (CpG at position 793, or CpG793 for short), are located between the TATA box and the transcription start site (TSS) (arrow in Figure 3C), consistent with previous reports suggesting functionally important CpG islands around the TSS^37^. CpG793 was among the CpGs that were eliminated in the construction of the low CpG pEF1s(low) promoter, consistent with the lower silencing rate observed for this promoter (Figure 2B).

To test for a functional role of CpG793, we mutated it to CC in pEF1s(orig) and in pEF1s(high). Conversely, we also reverted this position back to CG in pEF1s(low), where all 22 other CpGs including for CpG793 were mutated previously. Together, these constructs provided a set of controlled comparisons in which position 793 was either CC or CG in pEF1s(high), pEF1s(orig), and pEF1s(low) (Figure 3F, top).

We analyzed silencing rates (fraction per day normalized by remaining fraction, similar to Figure S3B) for each of these promoters. These rates were significantly reduced (in the “CC” variants of pEF1s(orig) (p<0.001) and pEF1s(low) (p<0.001), but not the pEF1s(high) promoters, compared to the CG variants (Figure 3F lower, and 3G). Further, silencing rates were similar between the CG variant of pEFs1(low) and the CC variant of pEFs1(orig) (barely significantly different with p=0.047), even though these two sequences systematically differed at 22 other CpGs. This indicates that the position 793 mutation could almost compensate for the combined effect of 22 other CpG mutations.

Finally, we asked if the observed differential silencing dynamics caused by the CC-CG mutation at position 793 could result from disruption or introduction of a known transcription factor binding site. We queried the surrounding sequence (±8 nt, 18 nts in total) against known mouse cis-regulartory elements in CIS-BP^38^, and filtered for hits expressed in CHO cells^39^ based on criteria suggested previously^40^ (Table S1). The only hits observed in both the “low” and “orig” promoters were Gmeb1 and Gmeb2, a pair of proteins that are involved in modulating glucocorticoid receptor-mediated transactivation.^41^ However, these proteins are not known to be directly involved in epigenetic regulation, to our knowledge. While we cannot rule out the possibility that the observed difference in silencing results from differential binding of sequence-dependent cis-factors, it is consistent with the explanation that methylation capability at this position has a disproportionate effect on silencing.

Together, we observed three key results: First, CpG methylation in promoters in the “still ON” population accumulated with time. Second, the silencing rate did not correlate with either methylation rate or total methylation, contrary to expectation. Third, we discovered a specific CpG position that plays a disproportionate, functional role in controlling silencing rate.

## CONCLUSIONS

While effects of sequence on DNMT-dependent gene silencing have long been observed, a controlled system in which to directly analyze the effects of sequence on silencing has not been available. Here, we constructed a library of synthetic promoters, featuring varying CpG content and DNMT-dependent silencing kinetics (Figure 1). Strikingly, silencing rate correlates directly with CpG content (Figure 2C). However, this correlation could not be explained by a corresponding effect of CpG content on methylation, as methylation accumulated at similar rates in all promoter variants (Figure 3B-3D). Finally, we observed evidence that a certain CpG (CpG793), located between the TATA box and the TSS, can play a disproportionate role in control of silencing rate (Figure 3F and 3G). Together, these results should provide a versatile set of components for engineering synthetic epigenetic circuits with desired silencing behaviors, as well as a foundation for future investigations of the mechanisms of DNMT-dependent silencing. Finally, our observation that the DNA sequence-based substrate of epigenetic modifications could alter the regulation dynamics might also apply into fully synthetic epigenetic circuits.^25^

A remaining mystery is why the rate of methylation accumulation is not correlated with the rate of silencing, nor with the CpG content of the promoter. Despite the lack of correlation between silencing rate and accumulated methylation, promoter silencing depended on the methylation activity of the recruited DNMT, as a catalytically inactive variant of DNMT3b was not able to initiate silencing in our system (Figure 2A), indicating that de novo DNA methylation is a necessary requirement for promoter silencing in this context.

A possible explanation could be that silencing requires at least two distinct steps, mediated by two types of cis-regulatory factors: the first binding methyl-CpG, and the second binding to CpG in a methylation-independent fashion. If only a small number of methyl-CpG are required for the first, methyl-dependent factor(s), then total CpG density could establish a rate-limiting step for advancing to a silent state. Examples of both types of proteins exist. Methyl-binding domain (MBD) proteins like MeCP2, MBD2 etc. are known to play key roles in methylation-dependent silencing.^6, 42^ At the same time, CpG islands are known to be able to initiate silencing by recruiting polycomb group proteins independent of methylation in embryonic stem cells differentiating into neurons.^43^ There are also “dual functional” proteins (e.g. TET1^44^ and KDM2B^45^) that bind to un-methylated CpGs but still promote gene silencing in some cell contexts. Further experiments could help to disentangle the roles of methylation-dependent and independent factors in controlling the rate of silencing.

Starting within two days after the release of dox, the active population remained in an actively expressing state (Figure 2A). DNA methylation is actively maintained, and thus unlikely to dilute out during this period without active recruitment of de-methylation enzymes.^46^ Therefore, the recruited DNMT3bCD protein may play an additional role in silencing beyond its catalytic activity as a methyltransferase. In fact, full length DNMT3b, even with its catalytic domain deactivated, has significant functions in epigenetic gene regulation through its protein and heterochromatin interacting domain.^35^ In this study, we specifically recruited the DNMT3b ”catalytic domain” (with the PWWP domain deleted). However, this protein still includes the ATRX domain that has been shown to associate with heterochromatin.^27^ Further investigation will be needed to identify the roles of methyltransferase-dependent and independent activities of DNMT3b.

One factor that could complicate our comparison between promoters is the distance from the recruited site (5xtetO) to the promoter’s core. This distance differed in promoters with additional inserts. However, the correlation of silencing rate with CpG density occurred among groups of constructs either lacking or containing the insert, when these groups were considered separately. This suggests that the change in distance to the core promoter (roughly 300 base pairs) in this system is not responsible for silencing rate correlation.

Additionally, our discovery of CpG793 playing a disproportionate role in determining silencing dynamics also suggested our model of correlation between CpG density and silencing rate is incomplete. A much larger set of promoter variants containing combinotory mutations on all CpGs may provide a more complete model accounting for the individual effects of each CpG. We believe this issue would be better resolved in the future using a massively parallel reporter assay approach that can access much larger numbers of promoters.

Finally, we note that phenotypically, our findings resemble the genetic mechanism of fragile X syndrome (FRX), in which an increased CGG repeat number upstream of the FRM1 gene’s promoter leads to hypermethylation and gene silencing during development.^47^ The exact molecular mechanism leading to silencing in FRX is not yet fully understood, but various hypotheses, including toxic secondary RNA structure^48^ and aberrant histone deacetylation^49^ have been proposed. It would be interesting to find out to what extent the mechanisms underlying the relationships observed here may be shared with those involved in FRX.

## MATERIALS AND METHODS

### Cell culture maintaining

CHO cells containing a human artificial chromosome (CHO-HAC)^29^ were cultured at 37°C, in a humidified atmosphere with 5% CO2. The growth media consisted of Alpha MEM Earle’s Salts (Irvine Scientific) with 10% Tet Approved FBS (Clontech Laboratories or Avantor) and 1X Penicillin/Streptomycin (Life Technologies) and 1x GlutaMax (Gibco) added. Cells are passaged according to the standard CHO-K1 cell (CCL-61, ATCC) procedure.

### Plasmid and cell line construction

All plasmids are constructed using standard cloning techniques, including Gibson Assembly (NEB) and GoldenGate Assembly (NEB). The plasmids and their maps are available for requests at Addgene (addgene.org/browse/article/28233817/).

The basal cell line expressing rTetR-DNMT3B (CD and CI version) is constructed by transfection and stable integration via the PiggyBAC system (System Biosciences), following manufacturer’s instructions, followed by blasticidin (Gibco) selection at 10 ug/ml for 5 days. The cells are then sorted for similar mCherry expression (Figure S1A), or single cloned further reporter integration (in the case of finer time course in Figure 2B). For integration of the reporter, methods similar to previous literature^18^ are used. Briefly, we co-transfected 600 ng reporter plasmid and 200 ng PhiC31 integrase plasmid using Lipofectamine 2000 (Invitrogen). After selection by geneticin (Gibco) at 400 ng/ml for 14 days (Supplementary S1B), cells were sorted (see below) to isolate the population with expression around the highest peak (Supplementary figure S1C, expected expression of single integration, as their system’s single integration rate should be close to 90% after selection^29^).

### Flow cytometry and fluorescence activated cell sorting (FACS)

Cells are washed by PBS, lifted by 0.25% EDTA-Trypsin (Gibco), and diluted in HBSS (Gibco) with 0.25% of BSA before flow cytometry. Flow cytometry experiments are performed either on MACSQuant VYB Analyzer (Miltenyi Biotec) or CytoFLEX (Beckman Coulter). Analysis of data is done with open source, in-house developed software, EasyFlow (https://github.com/AntebiLab/easyflow) or EasyFlowQ (https://github.com/ym3141/EasyFlowQ).

FACS is performed with SY3200 Cell Sorter (Sony) at Caltech FLow Cytometry Facility.

### Enzymatic methylation-specific sequencing and analysis

Cells were sorted as described above and immediately lysed for DNA extraction (DNeasy Blood & Tissue Kit, Qiagen). Total DNA is then converted with NEBNext Enzymatic Methyl-seq Conversion Module (NEB) according to the manufacturer’s instructions, and further amplified (EpiMark Hot Start Taq DNA Polymerase, NEB) with primers targeting a 2.5kb region containing TetO binding sites, promoter region and the gene body (nucleotide 1943-4438 on the none-pEF1s(orig) plasmid). The amplified targets are further prepared into library (Nextera XT Library Prep protocol, Illumina) and sequenced on the MiSeq (250bp pair ended, Illumina) platform.

The resulting reads were first trimmed and filtered by Trim Galore! (Babraham Institute) and then aligned and analyzed by Bismark^50^ and SAMtools^51^ to generate the methylation calling statistics.

### Data processing and statistical testing

For calculating the silenced fractions, the background silenced fraction (*S*_*dox*−_) from the no recruitment control sample (no dox) is subtracted from the observed silenced fraction (*S*_*dox*+_) from the with recruitment group experiment, and further normalized by the “fraction still available for silencing” (“still ON” fraction in the control 1 − *S*_*dox*−_. Consequently, the silenced rate of a given sample is calculated as 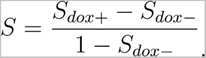.

All statistical testings in this study are Student’s t-test if not specified.

Error bars in Figure 3F and 3G are generated via bootstrapping. Specifically for each time point in Figure 3F and 3G, each of the three “with recruitment” samples are normalized to each of three “no recruitment” control samples, according to the method described above. Therefore a total of 9 data points are generated, and the error bars represent the standard deviation of these points.

### Data and code availability

Plasmids and their maps available for requests at Addgene (addgene.org/browse/article/28233817/). The key cell lines are available upon request. EM-Seq raw and processed data is deposited at Gene Expression Omnibus (GSE224403). Data and codes for analysis and generating figures are available at data.caltech (doi: 10.22002/ct5kt-cv878).

## SUPPORTING INFORMATION

### Supplementary table

**Supplementary Table S1:** Transcription factors that bind differentially only to the CG or CC (at CpG793) version of the promoters. Surrounding sequences (±8 nt, 18 nts in total) of CpG793 in three different promoters are queried against known mouse transcription factors (CIS-BP^38^) for potential differential TF binding between the CG and CC version. Potential hits are then compared in wild type CHO’s transcriptome data^39^, Genes that cannot be mapped in CHO transcriptome, or have a FKPM smaller than 1 are filtered out^40^. Among the hits, high expression genes (greater than 1/100 of the housekeeping Actb gene) are annotated with stars.

|  | pEF1s(low) | pEF1s(orig) | pEF1s(high) |
| --- | --- | --- | --- |
| CG variant | Tfe3, Tfeb, Mitf, Arntl, Mlx, Gmeb1, Gmeb2, Usf2* | Gmeb1, Gmeb2 | Dnmt1*, Kdm2b, Kmt2a |
| CC variant | Zbtb7bm | - | - |

### Supplementary figure and legends

**Figure S1.**
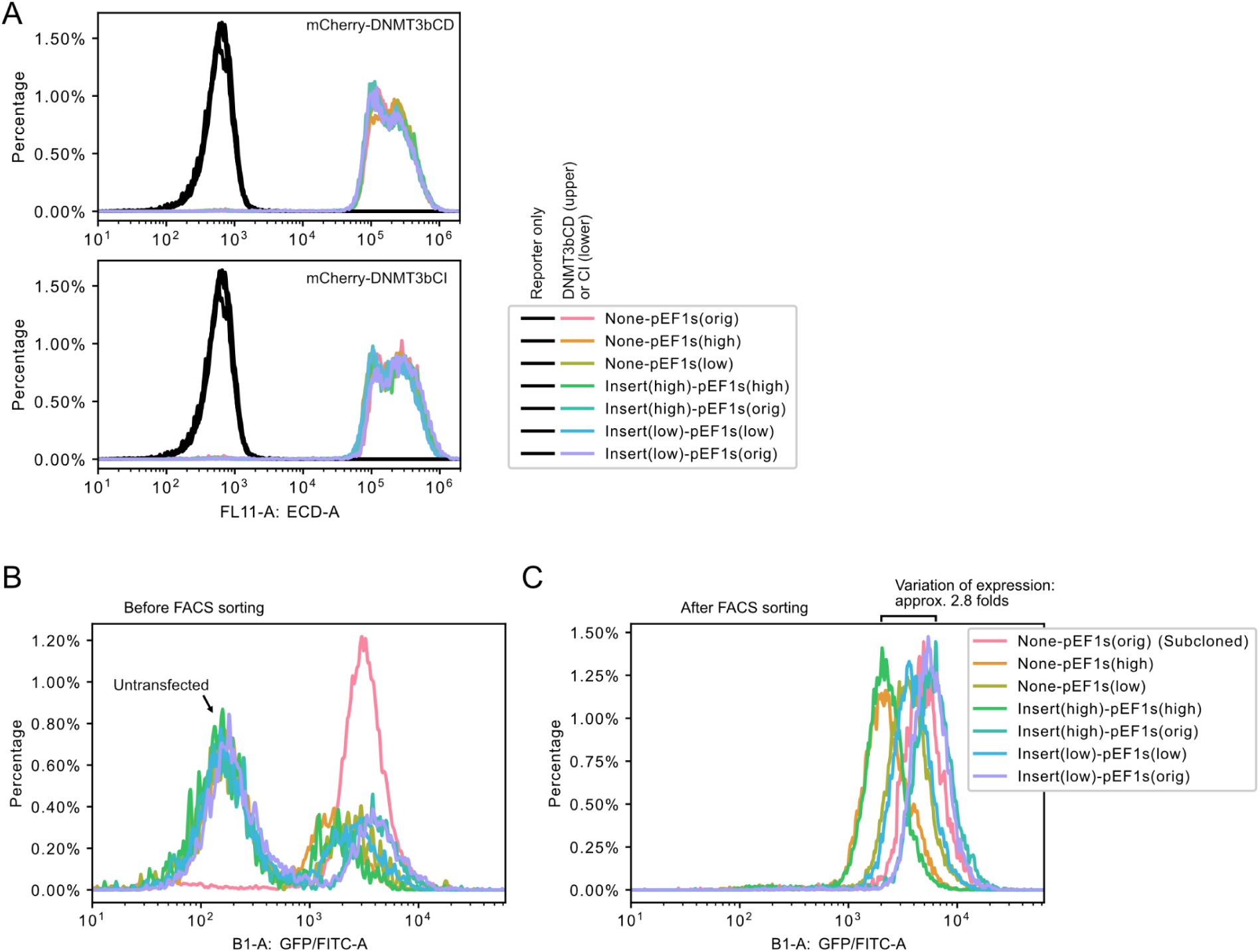
Generation of the cell line. A. Cell lines with different promoters have similar expression levels of DNMT3bCD (top) or DNMT3bCI (bottom). Cell lines that are transfected with DNMT3b (CD or CI) with co-expressing mCherry fluorescent protein, are selected by blasticidin and then sorted (see MATERIALS AND METHODS). All the lines have similar expression levels of DNMT3b (CD or CI). B-C. Site-specific integration of the reporters generates cell populations with uniform expression after FACS sorting: Cells are transfected and selected with geneticin for 14 days (B) (see MATERIALS AND METHODS). And then subcloned (None-pEF1s(orig)) or FACS sorted (all others) for further analysis (C).

**Figure S2.**
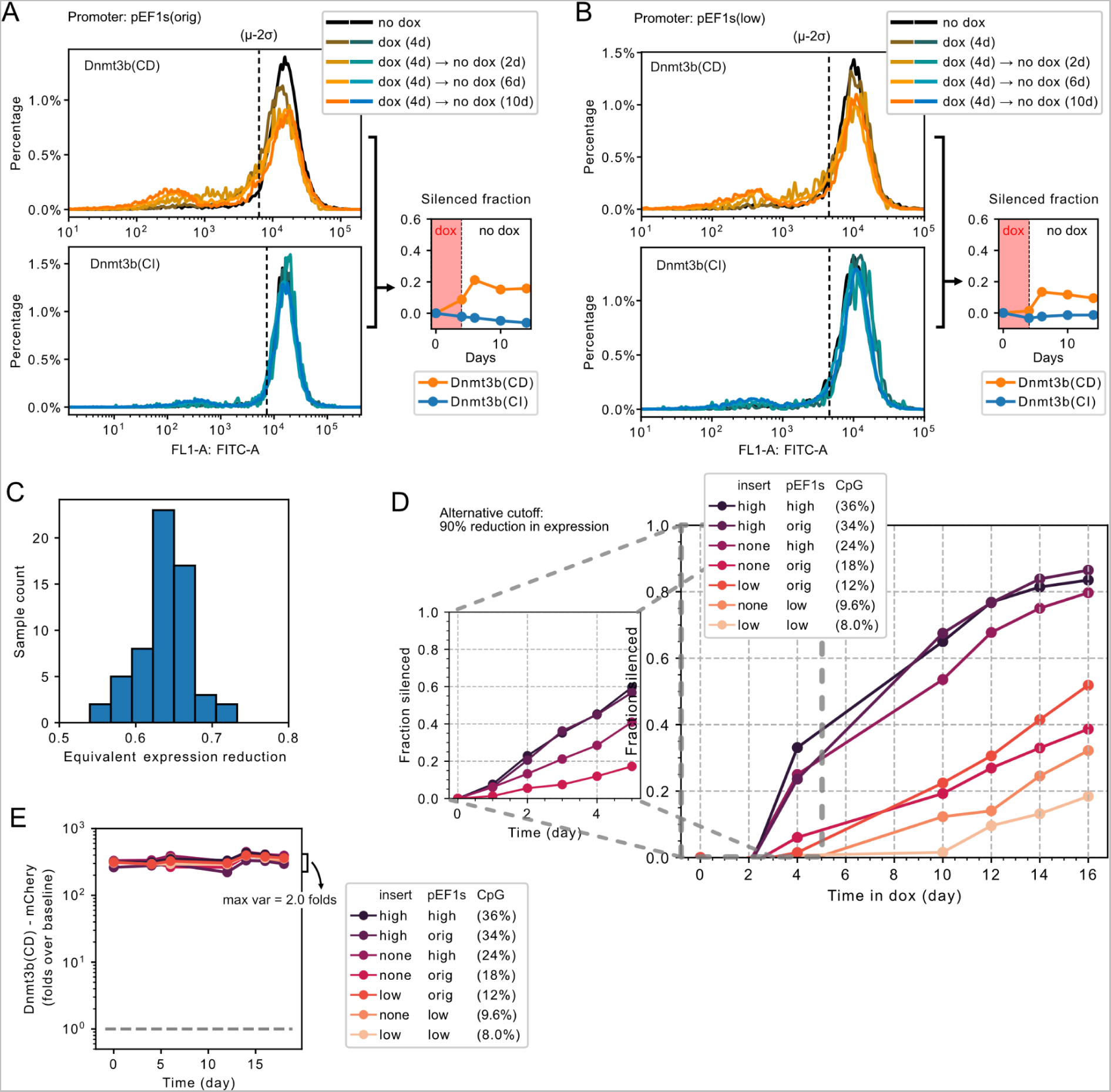
Promoters’ silencing rate correlates with their CpG content. A-B. Promoter silences with all-or-none kinetics when DNMT3bCD is recruited to the locus, and this silencing is dependent on DNMT3b’s catalytic activity. Same as Figure 2A except for promoter pEF1s(orig) (A) and pEF1s(low) (B). Lower silencing fractions are observed in general with these two promoters, but silenced populations remain stable after two days of dox release. DNMT3bCI does not silence either promoter. A. C. Comparison between the cutoff criteria we use (2σ from control expression, as in Figure 2 and Supplementary Figure S2A-B) to more traditional “expression reduction” criteria. Our criteria is equivalent to expression reduction from 54% to 71%, depending on the cell lines and time points, forming a single peaked distribution. B. D. Same time course as Figure 2B, with cutoff as 90% reduction in expression to determine silenced fraction. Though this method is less sensitive, similar correlation between promoters CpG density and silencing rates are observed. C. E. DNMT3bCD expression level maintained stably throughout the time course (Figure 2B). The mCherry co-expressed with DNMT3bCD maintained around 300 folds over the baseline throughout the time course, with maximal variation of 2 folds across different cell lines and time points.

**Figure S3:**
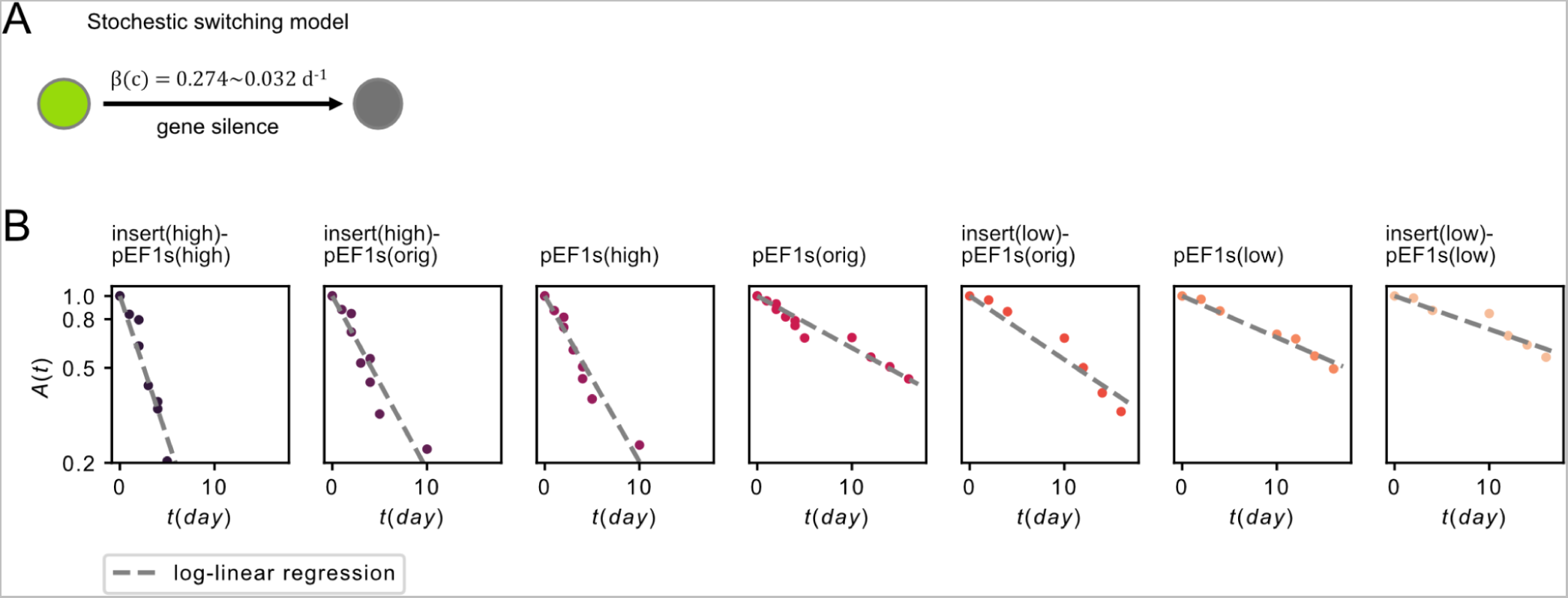
Promoters silencing dynamics follows A. Stochastic switching model of the silencing dynamics (see also Eq. 1). B. Time course from Figure 2B replotted as *A*(*t*) (1 - silenced fraction) against time . The *A*(*t*) (y-axis) is plotted in log scale as the model predicts a linear-log relationship. Regression is performed with the least-square method.

## AUTHOR INFORMATION

### Author contributions

Y.M.: Conceptualization, Formal analysis, Investigation, Writing; J.Z.: Formal analysis, Investigation; M.W.B.: Conceptualization, Supervision, Funding; M.B.E: Conceptualization, Supervision, Writing, Funding;

### Funding

This work is supported by the Defense Advanced Research Projects Agency under Contract No. HR0011-17-2-0008, by the National Institutes of Health grant RO1 HD075605A, and by National Science Foundation grant EF-2021552 under subaward UWSC10142. MBE is a Howard Hughes Medical Institute Investigator.

### Notes

M.W.B. is a founder and employee of Primordium Labs.

## ACKNOWLEDGEMENTS

We thank Jeff Park for technical assistance and advice; Rochelle Diamond and Jamie Tejirina at the Caltech Flow Cytometry and Cell Sorting Facility for technical advice and assistance. Yodai Takei, James Linton, Shiyu Xia and other members of the Elowitz lab for critical feedback on the manuscript; Lacramonica Bintu, and Matt Thomson for scientific input and advice.

This article is subject to HHMI’s Open Access to Publications policy. HHMI lab heads have previously granted a nonexclusive CC BY 4.0 license to the public and a sublicensable license to HHMI in their research articles. Pursuant to those licenses, the author-accepted manuscript of this article can be made freely available under a CC BY 4.0 license immediately upon publication.

## Notes

https://doi.org/10.22002/jdw94-3nh81

### Competing Interest Statement

The authors have declared no competing interest.

## REFERENCES

(1) Smith, Z. D.; Meissner, A. DNA Methylation: Roles in Mammalian Development. Nature Reviews Genetics. 2013, pp 204–220. https://doi.org/10.1038/nrg3354.

(2) Ehrlich, M. DNA Methylation in Cancer: Too Much, but Also Too Little. Oncogene 2002, 21 (35), 5400–5413.

(3) Greenberg, M. V. C.; Bourc’his, D. The Diverse Roles of DNA Methylation in Mammalian Development and Disease. Nat. Rev. Mol. Cell Biol. 2019, 20 (10), 590–607.

(4) McCabe, M. T.; Brandes, J. C.; Vertino, P. M. Cancer DNA Methylation: Molecular Mechanisms and Clinical Implications.Clinical Cancer Research. 2009, pp 3927–3937. https://doi.org/10.1158/1078-0432.ccr-08-2784.

(5) Attwood, J. T.; Yung, R. L.; Richardson, B. C. DNA Methylation and the Regulation of Gene Transcription. Cellular and Molecular Life Sciences CMLS. 2002, pp 241–257. https://doi.org/10.1007/s00018-002-8420-z.

(6) Zhu, H.; Wang, G.; Qian, J. Transcription Factors as Readers and Effectors of DNA Methylation. Nat. Rev. Genet. 2016,17 (9), 551–565.

(7) Lyko, F. The DNA Methyltransferase Family: A Versatile Toolkit for Epigenetic Regulation. Nature Reviews Genetics. 2018, pp 81–92. https://doi.org/10.1038/nrg.2017.80.

(8) Morita, S.; Noguchi, H.; Horii, T.; Nakabayashi, K.; Kimura, M.; Okamura, K.; Sakai, A.; Nakashima, H.; Hata, K.; Nakashima, K.; Hatada, I. Targeted DNA Demethylation in Vivo Using dCas9–peptide Repeat and scFv–TET1 Catalytic Domain Fusions. Nature Biotechnology. 2016, pp 1060–1065. https://doi.org/10.1038/nbt.3658.

(9) Choudhury, S. R.; Cui, Y.; Lubecka, K.; Stefanska, B.; Irudayaraj, J. CRISPR-dCas9 Mediated TET1 Targeting for Selective DNA Demethylation at BRCA1 Promoter. Oncotarget 2016, 7 (29), 46545–46556.

(10) Moore, L. D.; Le, T.;Fan, G. DNA Methylation and Its Basic Function. Neuropsychopharmacology 2013, 38 (1), 23–38.

(11) Lövkvist, C.; Dodd, I. B.; Sneppen, K.; Haerter, J. O. DNA Methylation in Human Epigenomes Depends on Local Topology of CpG Sites. Nucleic Acids Res. 2016, 44 (11), 5123–5132.

(12) Weber, M.; Hellmann, I.; Stadler, M. B.; Ramos, L.; Pääbo, S.; Rebhan, M.; Schübeler, D. Distribution, Silencing Potential and Evolutionary Impact of Promoter DNA Methylation in the Human Genome. Nat. Genet. 2007, 39 (4), 457–466.

(13) Haerter, J. O.; Lövkvist, C.; Dodd, I. B.; Sneppen, K. Collaboration between CpG Sites Is Needed for Stable Somatic Inheritance of DNA Methylation States. Nucleic Acids Res. 2014, 42 (4), 2235–2244.

(14) Bruno, S.; Williams, R. J.; Del Vecchio, D. Epigenetic Cell Memory: The Gene’s Inner Chromatin Modification Circuit. PLoS Comput. Biol. 2022, 18 (4), e1009961.

(15) Long, H. K.; King, H. W.; Patient, R. K.; Odom, D. T.; Klose, R. J. Protection of CpG Islands from DNA Methylation Is DNA-Encoded and Evolutionarily Conserved. Nucleic Acids Res. 2016, 44 (14), 6693–6706.

(16) Takahashi, Y.; Wu, J.; Suzuki, K.; Martinez-Redondo, P.; Li, M.; Liao, H.-K.; Wu, M.-Z.; Hernández-Benítez, R.; Hishida, T.; Shokhirev, M. N.; Esteban, C. R.; Sancho-Martinez, I.; Belmonte, J. C. I. Integration of CpG-Free DNA Induces de Novo Methylation of CpG Islands in Pluripotent Stem Cells. Science 2017, 356 (6337), 503–508.

(17) Singer, Z. S.; Yong, J.; Tischler, J.; Hackett, J. A.; Altinok, A.; Surani, M. A.; Cai, L.; Elowitz, M. B. Dynamic Heterogeneity and DNA Methylation in Embryonic Stem Cells. Mol. Cell 2014, 55 (2), 319–331.

(18) Bintu, L.; Yong, J.; Antebi, Y. E.; McCue, K.; Kazuki, Y.; Uno, N.; Oshimura, M.; Elowitz, M. B. Dynamics of Epigenetic Regulation at the Single-Cell Level. Science 2016, 351 (6274), 720–724.

(19) Ziller, M. J.; Gu, H.; Müller, F.; Donaghey, J.; Tsai, L. T.-Y.; Kohlbacher, O.; De Jager, P. L.; Rosen, E. D.; Bennett, D. A.; Bernstein, B. E.; Gnirke, A.; Meissner, A. Charting a Dynamic DNA Methylation Landscape of the Human Genome. Nature 2013, 500 (7463), 477–481.

(20) Kungulovski, G.; Jeltsch, A. Epigenome Editing: State of the Art, Concepts, and Perspectives. Trends Genet. 2016, 32 (2), 101–113.

(21) Nakamura, M.; Gao, Y.; Dominguez, A. A.; Qi, L. S. CRISPR Technologies for Precise Epigenome Editing. Nature Cell Biology. 2021, pp 11–22. https://doi.org/10.1038/s41556-020-00620-7.

(22) Van, M. V.; Fujimori, T.; Bintu, L. Nanobody-Mediated Control of Gene Expression and Epigenetic Memory. Nat. Commun. 2021, 12 (1), 537.

(23) Liu, X. S.; Shawn Liu, X.; Wu, H.; Ji, X.; Stelzer, Y.; Wu, X.; Czauderna, S.; Shu, J.; Dadon, D.; Young, R. A.; Jaenisch, R. Editing DNA Methylation in the Mammalian Genome. Cell. 2016, pp 233–247.e17. https://doi.org/10.1016/j.cell.2016.08.056.

(24) Nuñez, J. K.; Chen, J.; Pommier, G. C.; Cogan, J. Z.; Replogle, J. M.; Adriaens, C.; Ramadoss, G. N.; Shi, Q.; Hung, K. L.; Samelson, A. J.; Pogson, A. N.; Kim, J. Y. S.; Chung, A.; Leonetti, M. D.; Chang, H. Y.; Kampmann, M.; Bernstein, B. E.; Hovestadt, V.; Gilbert, L. A.; Weissman, J. S. Genome-Wide Programmable Transcriptional Memory by CRISPR-Based Epigenome Editing. Cell 2021, 184 (9), 2503–2519.e17.

(25) Park, M.; Patel, N.; Keung, A. J.; Khalil, A. S. Engineering Epigenetic Regulation Using Synthetic Read-Write Modules. Cell 2019, 176 (1-2), 227–238.e20.

(26) Urlinger, S.; Baron, U.; Thellmann, M.; Hasan, M. T.; Bujard, H.; Hillen, W. Exploring the Sequence Space for Tetracycline-Dependent Transcriptional Activators: Novel Mutations Yield Expanded Range and Sensitivity. Proc. Natl. Acad. Sci. U. S. A. 2000, 97 (14), 7963–7968.

(27) Chen, T.; Tsujimoto, N.; Li, E. The PWWP Domain of Dnmt3a and Dnmt3b Is Required for Directing DNA Methylation to the Major Satellite Repeats at Pericentric Heterochromatin. Mol. Cell. Biol. 2004, 24 (20), 9048–9058.

(28) Zacharias, D. A.; Violin, J. D.; Newton, A. C.; Tsien, R. Y. Partitioning of Lipid-Modified Monomeric GFPs into Membrane Microdomains of Live Cells. Science 2002, 296 (5569), 913–916.

(29) Yamaguchi, S.; Kazuki, Y.; Nakayama, Y.; Nanba, E.; Oshimura, M.; Ohbayashi, T. A Method for Producing Transgenic Cells Using a Multi-Integrase System on a Human Artificial Chromosome Vector. PLoS One 2011, 6 (2), e17267.

(30) Argentova, V. V.; Aliev, T. K.; Toporova, V. A.; Rybchenko, V. S.; Dolgikh, D. A.; Kirpichnikov, M. P. Studies on the Influence of Different Designs of Eukaryotic Vectors on the Expression of Recombinant IgA. Moscow University Biological Sciences Bulletin. 2017, pp 63–68. https://doi.org/10.3103/s0096392517020018.

(31) Oliveira, D. S. L. de; Paredes, V.; Caixeta, A. V.; Henriques, N. M.; Wear, M. P.; Albuquerque, P.; Felipe, M. S. S.; Casadevall, A.; Nicola, A. M. Hinge Influences in Murine IgG Binding to Cryptococcus Neoformans Capsule. Immunology 2022, 165 (1), 110–121.

(32) Fu, X.; Zhu, J.; Duan, Y.; Lu, P.; Zhang, K. CRISPR/Cas9 Mediated Somatic Gene Therapy for Insertional Mutations: The Mouse Model. Precis Clin Med 2021, 4 (3), 168–175.

(33) Sayers, E. W.; Bolton, E. E.; Brister, J. R.; Canese, K.; Chan, J.; Comeau, D. C.; Connor, R.; Funk, K.; Kelly, C.; Kim, S.; Madej, T.; Marchler-Bauer, A.; Lanczycki, C.; Lathrop, S.; Lu, Z.; Thibaud-Nissen, F.; Murphy, T.; Phan, L.; Skripchenko, Y.; Tse, T.; Wang, J.; Williams, R.; Trawick, B. W.; Pruitt, K. D.; Sherry, S. T. Database Resources of the National Center for Biotechnology Information. Nucleic Acids Res. 2022, 50 (D1), D20–D26.

(34) Jia, L.; Mao, Y.; Ji, Q.; Dersh, D.; Yewdell, J. W.; Qian, S.-B. Decoding mRNA Translatability and Stability from the 5’ UTR. Nat. Struct. Mol. Biol. 2020, 27 (9), 814–821.

(35) Nowialis, P.; Lopusna, K.; Opavska, J.; Haney, S. L.; Abraham, A.; Sheng, P.; Riva, A.; Natarajan, A.; Guryanova, O.; Simpson, M.; Hlady, R.; Xie, M.; Opavsky, R. Catalytically Inactive Dnmt3b Rescues Mouse Embryonic Development by Accessory and Repressive Functions. Nat. Commun. 2019, 10 (1), 4374.

(36) Vaisvila, R.; Ponnaluri, V. K. C.; Sun, Z.; Langhorst, B. W.; Saleh, L.; Guan, S.; Dai, N.; Campbell, M. A.; Sexton, B. S.; Marks, K.; Samaranayake, M.; Samuelson, J. C.; Church, H. E.; Tamanaha, E.; Corrêa, I. R., Jr; Pradhan, S.; Dimalanta, E. T.; Evans, T. C., Jr; Williams, L.; Davis, T. B. Enzymatic Methyl Sequencing Detects DNA Methylation at Single-Base Resolution from Picograms of DNA. Genome Res. 2021, 31 (7), 1280–1289.

(37) Fenouil, R.; Cauchy, P.; Koch, F.; Descostes, N.; Cabeza, J. Z.; Innocenti, C.; Ferrier, P.; Spicuglia, S.; Gut, M.; Gut, I.; Andrau, J.-C. CpG Islands and GC Content Dictate Nucleosome Depletion in a Transcription-Independent Manner at Mammalian Promoters. Genome Research. 2012, pp 2399–2408. https://doi.org/10.1101/gr.138776.112.

(38) Weirauch, M. T.; Yang, A.; Albu, M.; Cote, A. G.; Montenegro-Montero, A.; Drewe, P.; Najafabadi, H. S.; Lambert, S. A.; Mann, I.; Cook, K.; Zheng, H.; Goity, A.; van Bakel, H.; Lozano, J.-C.; Galli, M.; Lewsey, M. G.; Huang, E.; Mukherjee, T.; Chen, X.; Reece-Hoyes, J. S.; Govindarajan, S.; Shaulsky, G.; Walhout, A. J. M.; Bouget, F.-Y.; Ratsch, G.; Larrondo, L. F.; Ecker, J. R.; Hughes, T. R. Determination and Inference of Eukaryotic Transcription Factor Sequence Specificity. Cell 2014, 158 (6), 1431–1443.

(39) Kondratova, A.; Watanabe, T.; Marotta, M.; Cannon, M.; Segall, A. M.; Serre, D.; Tanaka, H. Replication Fork Integrity and Intra-S Phase Checkpoint Suppress Gene Amplification. Nucleic Acids Res. 2015, 43 (5), 2678–2690.

(40) Nagaraj, N.; Wisniewski, J. R.; Geiger, T.; Cox, J.; Kircher, M.; Kelso, J.; Pääbo, S.; Mann, M. Deep Proteome and Transcriptome Mapping of a Human Cancer Cell Line. Mol. Syst. Biol. 2011, 7, 548.

(41) Zeng, H.; Kaul, S.; Simons, S. S., Jr. Genomic Organization of Human GMEB-1 and Rat GMEB-2: Structural Conservation of Two Multifunctional Proteins. Nucleic Acids Res. 2000, 28 (8), 1819–1829.

(42) Spruijt, C. G.; Vermeulen, M. DNA Methylation: Old Dog, New Tricks? Nat. Struct. Mol. Biol. 2014, 21 (11), 949–954.

(43) Deaton, A. M.; Bird, A. CpG Islands and the Regulation of Transcription. Genes Dev. 2011, 25 (10), 1010–1022.

(44) Wu, H.; D’Alessio, A. C.; Ito, S.; Xia, K.; Wang, Z.; Cui, K.; Zhao, K.; Sun, Y. E.; Zhang, Y. Dual Functions of Tet1 in Transcriptional Regulation in Mouse Embryonic Stem Cells. Nature 2011, 473 (7347), 389–393.

(45) Farcas, A. M.; Blackledge, N. P.; Sudbery, I.; Long, H. K.; McGouran, J. F.; Rose, N. R.; Lee, S.; Sims, D.; Cerase, A.; Sheahan, T. W.; Koseki, H.; Brockdorff, N.; Ponting, C. P.; Kessler, B. M.; Klose, R. J. KDM2B Links the Polycomb Repressive Complex 1 (PRC1) to Recognition of CpG Islands. Elife 2012, 1, e00205.

(46) Maeder, M. L.; Angstman, J. F.; Richardson, M. E.; Linder, S. J.; Cascio, V. M.; Tsai, S. Q.; Ho, Q. H.; Sander, J. D.; Reyon, D.; Bernstein, B. E.; Costello, J. F.; Wilkinson, M. F.; Joung, J. K. Targeted DNA Demethylation and Activation of Endogenous Genes Using Programmable TALE-TET1 Fusion Proteins. Nat. Biotechnol. 2013, 31 (12), 1137–1142.

(47) Garber, K. B.; Visootsak, J.; Warren, S. T. Fragile X Syndrome. Eur. J. Hum. Genet. 2008, 16 (6), 666–672.

(48) Ajjugal, Y.; Kolimi, N.; Rathinavelan, T. Secondary Structural Choice of DNA and RNA Associated with CGG/CCG Trinucleotide Repeat Expansion Rationalizes the RNA Misprocessing in FXTAS. Sci. Rep. 2021, 11 (1), 8163.

(49) Coffee, B.; Zhang, F.; Warren, S. T.; Reines, D. Acetylated Histones Are Associated with FMR1 in Normal but Not Fragile X-Syndrome Cells. Nat. Genet. 1999, 22 (1), 98–101.

(50) Krueger, F.; Andrews, S. R. Bismark: A Flexible Aligner and Methylation Caller for Bisulfite-Seq Applications. Bioinformatics. 2011, pp 1571–1572. https://doi.org/10.1093/bioinformatics/btr167.

(51) Danecek, P.; Bonfield, J. K.; Liddle, J.; Marshall, J.; Ohan, V.; Pollard, M. O.; Whitwham, A.; Keane, T.; McCarthy, S. A.; Davies, R. M.; Li, H. Twelve Years of SAMtools and BCFtools. Gigascience 2021, 10 (2). https://doi.org/10.1093/gigascience/giab008.

